# Rapid carbon accumulation at a saltmarsh restored by managed realignment far exceeds carbon emitted in site construction

**DOI:** 10.1101/2021.10.12.464124

**Authors:** Hannah L. Mossman, Nigel Pontee, Katie Born, Peter J. Lawrence, Stuart Rae, James Scott, Beatriz Serato, Robert B. Sparkes, Martin J.P. Sullivan, Rachel M. Dunk

## Abstract

Increasing attention is being paid to the carbon sequestration and storage services provided by coastal blue carbon ecosystems such as saltmarshes. Sites restored by managed realignment, where existing sea walls are breached to reinstate tidal inundation to the land behind, have considerable potential to accumulate carbon through deposition of sediment brought in by the tide and burial of vegetation in the site. While this potential has been recognised, it is not yet a common motivating factor for saltmarsh restoration, partly due to uncertainties about the rate of carbon accumulation and how this balances against the greenhouse gases emitted during site construction. We use a combination of field measurements over four years and remote sensing to quantify carbon accumulation at a large managed realignment site, Steart Marshes, UK. Sediment accumulated rapidly at Steart Marshes (mean of 75 mm yr^-1^) and had a high carbon content (4.4% total carbon, 2.2% total organic carbon), resulting in carbon accumulation of 36.6 t ha^-1^ yr^-1^ total carbon (19.4 t ha^- 1^ yr^-1^ total organic carbon). This rate of carbon accumulation is an order of magnitude higher than reported in many other restored saltmarshes, and is higher although more similar to values previously reported from another hypertidal system (Bay of Fundy, Canada). The estimated carbon emissions associated with the construction of the site were ∼2-4% of the observed carbon accumulation during the study period, supporting the view that managed realignment projects in such settings are likely to have significant carbon accumulation benefits. We outline further considerations that are needed to move towards a full carbon budget for saltmarsh restoration.

## Introduction

Earth’s ecosystems exhibit overall net carbon uptake, causing increases in atmospheric CO_2_ to be smaller than expected from fossil emissions and land-use change [1]. They also contain substantial carbon stocks, which currently store carbon out of the atmosphere but are sensitive to changes in climate or land-use [2, 3]. Coastal ‘blue carbon’ ecosystems, including saltmarshes, are especially carbon dense and sequester carbon at a rate an order of magnitude faster than terrestrial ecosystems [4]. Both allochthonous and autochthonous carbon is sequestered in saltmarshes; carbon accumulates in saltmarshes as sediment carried in by the tide is deposited, and this sediment buries saltmarsh plant remains. Globally, the ∼5.5 million hectares of saltmarshes [5] are estimated to accumulate carbon at an average rate of ∼2.4 t C ha^-1^ yr^-1^ [6]. Despite their importance, ∼50% of saltmarsh area has been lost, particularly through reclamation for agriculture or urbanisation, or degraded [7], with annual losses of 1-2% [8, 9].

In response to losses of saltmarsh and its associated biodiversity, ‘no net loss’ policies have sought to protect remaining wetlands and create new habitat [10], contributing to over 100,000 ha of intertidal wetland creation over the last 30 years [11]. However, the pace of global wetland creation is not sufficient to offset losses, where a key barrier is the availability of project financing [12].Payments for ecosystem services, such as flood protection or biodiversity, offer potential financial mechanisms for saltmarsh creation or restoration [13]. Carbon accumulation (and thus climate mitigation) has been recognised as a potential benefit of saltmarsh restoration, and could therefore provide a further motivation for site creation [14, 15].

Robustly quantifying the rate of carbon accumulation on restored saltmarshes will be necessary if carbon finance mechanisms are to be developed [16] and is also important to enable saltmarsh restoration to be properly included in national carbon budgets [17]. Furthermore, rising sea levels threaten existing saltmarshes, and the climate sensitivity of their carbon stocks and fluxes needs to be quantified [18]. While saltmarsh restoration could potentially compensate for loss of natural saltmarshes, given known differences in topography and ecology [19, 20], it may not be appropriate to assume that restored or created marshes will ultimately store carbon at a rate comparable to natural saltmarshes [21]. Thus it is also important to determine any differences between carbon accumulation and sequestration in natural and restored saltmarsh.

Previous attempts to quantify actual or potential carbon accumulation following saltmarsh restoration have used a variety of techniques: (a) spatially explicit models to predict landscape-scale carbon accumulation based on observed carbon accumulation in natural habitats [22]; (b) measurements at a single time-point to take a snapshot of carbon stocks [23]; (c) restored saltmarshes of different ages as a space-for-time substitution to estimate the rate of carbon accumulation [24]; and (d) repeat measurements of the elevation of sediment surface to quantify sediment deposition rates [25]. While all approaches highlight the potential for saltmarsh restoration to lead to carbon accumulation, each has limitations when used in isolation. A further challenge is that previous studies have either assessed only total carbon (which does not distinguish organic carbon from inorganic carbon such as biogenic or lithogenic carbonates), or have quantified organic carbon using loss on ignition, which is known to have poor accuracy and large uncertainties [26].

A further consideration when evaluating the net carbon benefit of a saltmarsh restoration or creation project is the balance between the carbon costs of constructing the site (e.g. building new flood defences inland and breaching the existing embankments, termed “managed realignment”) and the carbon accumulation provided by the site [e.g. 27]. If project carbon costs are high relative to the rate of carbon accumulation, it may take years for the site to pay off the debt of construction [28].

This research aims to evaluate carbon costs and benefits from saltmarsh creation through managed realignment, using a novel combination of techniques. Over the course of several annual cycles we use remote sensing, field measurements and robust laboratory techniques to quantify total and organic carbon accumulation in an evolving saltmarsh in the first years after restoration. This allows us to reliably quantify the amount and rate of carbon accumulation following restoration. We then assess the carbon emissions incurred during site construction before identifying additional requirements for producing a full carbon budget for saltmarsh restoration.

## Materials and Methods

### Study site

Steart Marshes (Somerset, UK; 51.20 N, 3.05 W) is a 250-ha managed realignment site, forming part of a larger 400 ha complex of restored wetland habitats managed by the Wildfowl and Wetlands Trust. It was constructed to create new intertidal habitat in compensation for previous losses, and to provide enhanced flood defences [29]. Prior to site construction, the land was under a mix of agricultural uses, including permanent pasture (i.e. pasture had been the land use over many years), grass ley (part of cyclical arable land management) and arable (winter wheat, barley, oilseed rape and maize) (Fig. 1a). The site lies near the mouth of the River Parrett which drains a catchment of interbedded limestone and mudstone [30] and flows into the Severn Estuary. Hydrodynamic processes in the Parrett are dominated by a large tidal range which gives rise to strong tidal flows and large intertidal areas. At Hinkley, just to the west of the Parrett Estuary mouth, the mean spring tides have a high water height of 5.6 mODN and a low water height of −5.1 mODN, giving a range of approximately 11m [31, 32].

**Figure 1.**
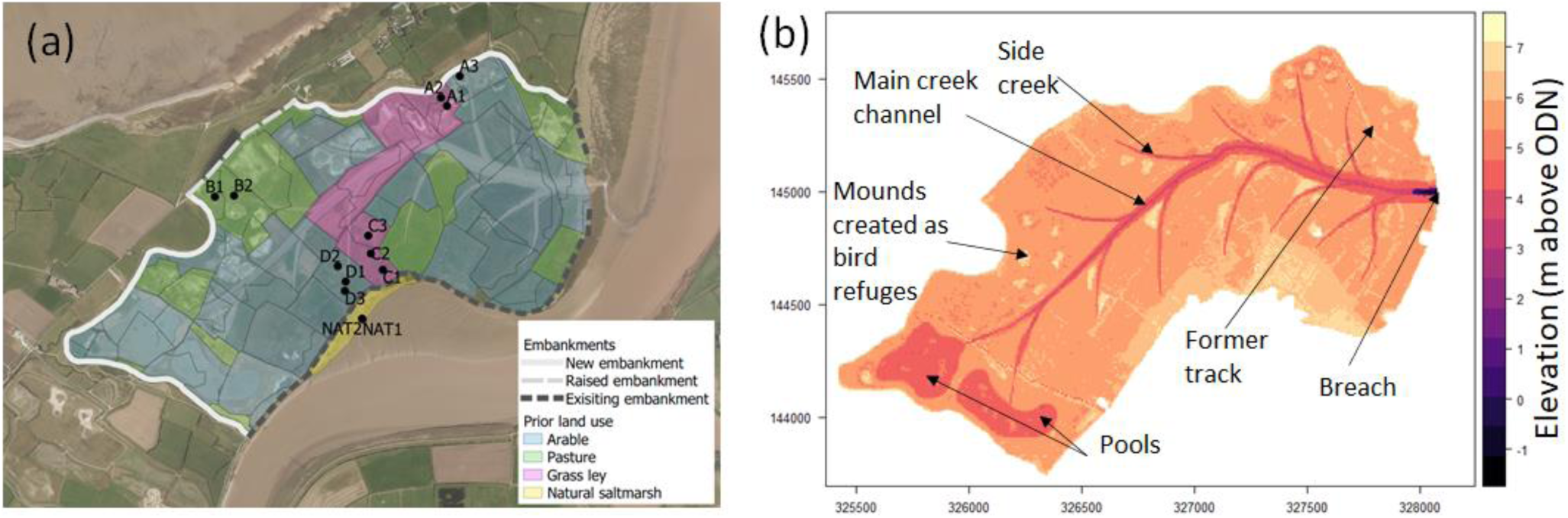
Design and construction elements of Steart managed realignment, Somerset, UK. a) Land use prior to the start of site construction in 2012, and locations of sampling points and the flood embankments constructed (new) or modified (raised) during the project; existing embankments that remained after the project are also shown. Land use was derived from Centre for Ecology and Hydrology Land Cover Map 2007 [66] and the project environmental statement [67]. Base aerial image from 2014 [68]. b) Elevations across the site showing design and location of creek network, lagoons and islands. The location of the breach is also shown. Elevations based on LiDAR data from October 2014 [36].

The construction of the managed realignment site started in early 2012, comprising the excavation of a creek network and pools, the construction of new flood defence embankments and the raising of a small length of existing embankment. The creek network (7.6 km total length) was designed to meet the geomorphological requirements of the scheme (see [33] for details), aid establishment of intertidal habitat, and minimise material transport distances by enabling construction of the required embankments from the excavated material [29]. In total, 4.75 km of new 4 m high or raised flood defence embankments were constructed (Fig. 1a). All material used in the construction of the new embankments was obtained from the site, i.e. embankments were created from clays excavated from within site and no concrete was used in embankment construction. Several lagoons were excavated to enhance habitat provision for birds and fish, and islands were created from excess material to provide protected roosting and nesting locations for birds at elevations high enough to avoid excessive inundation by the tide [29]. In total, 489,422 m^3^ of material was excavated and moved within the site during construction. A single, 250 m wide breach in the sea wall was created in September 2014, allowing regular tidal inundation to occur (further details of the breach are provided in [31]).

### Field sampling design

Four areas of the site were selected for regular sampling, first an area substantially disturbed by earth moving vehicles during construction (Site A, Fig. 1a) and three sites based on prior land use, permanent pasture (Site B), grass ley (Site C) and arable (Site D). Within these areas, we selected three sampling locations, stratified by the elevation prior to restoration of tidal inundation; the area of permanent pasture was relatively homogenous in elevation and so we only selected two sampling sites. To act as a natural reference, we selected a neighbouring area of pioneer saltmarsh (mostly bare ground with some *Spartina anglica*) and an area of saltmarsh with plant communities similar to those anticipated to establish on the managed realignment site, i.e. those dominated by *Puccinellia maritima* and *Aster tripolium* (NAT, Fig. 1a). This gave a total of thirteen regular sampling locations within five sampling sites.

### Sediment collection, preparation and storage

Sediments were sampled at each location immediately prior to restoration (28 August 2014, Sites A-D but not natural marsh), in December 2014 and then once or twice annually in 2015, 2016 and 2017, giving one pre-restoration and six post-restoration sampling time points (see Table S1 for full details). Cores of 30-50cm were collected using a soil auger and sectioned into 5-10 cm lengths for later analysis. In total, we collected 78 cores, resulting in 596 samples. Depending on site conditions, surface silts deposited post-breach were sometimes difficult to sample using a soil auger as they were either prone to compression or highly friable. In these cases, we collected undisturbed surface samples using adapted syringe tubes and/or collected the full surface sediment plates (down to base of mud cracks), and then sampled deeper sediments by taking a core between mud cracks. The horizon between the deposited silts and the underlying agricultural soils was determined through visual inspection of the cores (prior to sub-sampling) and the depth (in core and from surface) was recorded. The horizon was readily identifiable through a change in colour and texture of the soils, and by the presence of remnant vegetation and roots. Samples with a defined volume of 5 cm^3^ were taken from the above-horizon section of the core or directly from surface sediments for dry bulk density measurements [34].

All samples were stored at 4°C prior to analysis. Dry bulk density was determined by drying the samples of a known volume to a constant weight at 105°C. The remaining core samples were dried at 60°C, covered, in aluminium trays/glass jars for approximately 96 hours, then ground using a pestle and mortar to ensure a homogeneous sample for further analysis.

### Quantifying carbon composition of the sediment

We quantified the total carbon (TC) in all core sections collected. Total carbon contents were measured on dried, ground sediment samples using elemental analysis (LECO CR-412 Carbon Analyser (LECO Corporation, MI, USA) and Vario EL Cube (Elementar, Germany) instruments). Certified Reference Materials were analysed on both instruments and replicates of an internal standard (bulk sample of surface sediments collected from the site) were included in all instrument runs. The measured carbon content of the CRM on the LECO instrument was consistently higher than the certified value (Leco Soil Standard 502-062 (n=42), measured %C = 2.12±0.02, certified %C = 2.01±0.03), while analysis of the CRM on the Elementar instrument showed excellent agreement (Elemental Microanalysis Ltd Soil Standard B2184 (n=6), measured %C = 2.29±0.06, certified %C = 2.31±0.06). Accordingly, all LECO measurements were multiplied by a correction factor of 0.95(2.01/2.12).The corrected analysis of the internal standard on the LECO instrument showed excellent agreement with the Elementar analysis (LECO (n=83), %C = 2.74±0.10; Elementar (n=18), %C = 2.68±0.11)..

To quantify total organic carbon (TOC), we selected one core from each site (A-D and NAT) from the most recent sampling period; for the restored marsh (sites A-D), TOC was quantified on the newly accreted sediment (above horizon) only. These samples underwent acid digestion to remove inorganic carbon. Excess 1N HCl was added, and the samples were placed on a hotplate for three hours at 80 °C [35]. Following the acid digest, the supernatant liquor was decanted, and the samples rinsed 3 times with deionised water before being taken to dryness. This process removes calcium and magnesium carbonates (aragonite, calcite, and dolomite), along with other water- or acid-soluble minerals, whilst minimising loss of labile organic matter. Decarbonated samples were analysed on the Elementar instrument. Sample mass was recorded before and after decarbonation, with TOC values corrected to original sample mass.

### Quantifying sediment deposition and erosion

Multiple Digital Terrain Models (DTM) at 50 cm horizontal resolution were obtained for the site, derived from airborne LiDAR data [36]. The final pre-breach imagery, from 10 July 2014, pre-dates significant earth movement on site and is therefore unsuitable for use as a baseline. Instead, we have used the first post-breach imagery, from 31 October 2014 (57 days post breach), as a baseline for sediment accumulation on site. Between the date of the breach (4 September 2014) and the date of the imagery used, approximately 37 tides overtopped the creek banks and flooded some of the marsh surface (tides greater than 5.7 m ODN at the nearest available tide gauge, Hinkley Point (data from UK National Tide Gauge Network)). We obtained LiDAR DTMs for eleven further time points after breaching (see Table 1). Downloaded DTMs were processed in Rv4.02 [37] using the “raster” package [38]. Tiles were merged before being clipped by the site area. The site area was defined by manually drawing a polygon around the crest of the flood embankment to remove areas outside of the site. We then restricted analyses to locations subject to tidal inundation which were taken to be those areas below 7.07 m ODN, which is the level of the highest astronomical tides at the nearest port, Burnham-on-Sea [32]. The first DTM available after the breach (31 October 2014) was clipped to locations below 7.07 m and the resulting polygon (with an area of 244.7 ha) used to clip the remaining DTMs.

**Table 1.** Sedimentation at Steart Marshes measured by comparing Lidar DTMs to a baseline survey on 31 October 2014, 57 days after sea defences were breached.

| Survey date | Days since breach | Sedimentation rate (m yr<sup>-1</sup>) |  | Mean sediment depth (m) | Cumulative sediment volume (m<sup>3</sup>) | Cumulative carbon (95% confidence intervals) (t) | Cumulative organic carbon (95% confidence intervals) (t) |
| --- | --- | --- | --- | --- | --- | --- | --- |
|  |  | since start* | since previous survey |  |  |  |  |
| 24/01/2015 | 142 | 0.449 | 0.449 | 0.104 | 255646 | 12338<br>(2620-25697) | 6533<br>(1308-14101) |
| 05/04/2015 | 213 | 0.258 | 0.029 | 0.110 | 269282 | 13050<br>(3178-26582) | 6910<br>(1617-14704) |
| 04/06/2015 | 273 | 0.217 | 0.112 | 0.128 | 314126 | 15186<br>(4750-29578) | 8041<br>(2384-16392) |
| 31/07/2015 | 330 | 0.127 | -0.216 | 0.095 | 231555 | 11271<br>(1708-24269) | 5968<br>(875-13455) |
| 28/09/2015 | 389 | 0.153 | 0.277 | 0.139 | 341190 | 16491<br>(5625-31303) | 8732<br>(2837-17431) |
| 07/04/2016 | 581 | 0.110 | 0.034 | 0.157 | 384983 | 18626<br>(7016-34395) | 9863<br>(3536-19060) |
| 02/03/2017 | 910 | 0.092 | 0.064 | 0.215 | 525227 | 25507<br>(11315-44184) | 13506<br>(5614-24670) |
| 06/10/2017 | 1128 | 0.067 | -0.028 | 0.198 | 483641 | 23490<br>(10045-41212) | 12438<br>(5004-23134) |
| 01/04/2018 | 1305 | 0.073 | 0.105 | 0.249 | 608657 | 29541<br>(13545-49886) | 15642<br>(6733-28069) |
| 13/09/2018 | 1470 | 0.075 | 0.096 | 0.292 | 714513 | 34642<br>(16398-57400) | 18343<br>(8090-32402) |

Filtered DTM data should represent the ground elevations, but filtering does not completely remove dense, relatively short vegetation. Vegetation cover at the site in the first three years was sparse (Fig. S1, H Mossman pers. obs.) and so we do not consider this an issue for those years; in the latest year, vegetation cover was denser and extensive, but unvegetated areas remained (H Mossman pers. obs.). The 50 cm resolution cannot account for surface morphology smaller than this (e.g. surface desiccation cracking, which was observed during summer months). We also observed sediment dewatering and shrinkage during dry periods, but these changes were small compared to interannual changes in elevation (Table 1). Pontee and Serato [31] quantified the variation in elevation of control points between years (the same LiDAR datasets we use) and found a mean vertical error of ±0.04 m.

Changes in elevation were calculated between each time point by subtracting the DTM of the first time point from the DTM of the more recent time point. Cumulative changes in elevation were calculated relative to the first post-breach DTM (31 October 2014, 57 days after breach). We calculated the mean elevation change across raster pixels, which was then converted to total change in sediment volume by multiplying by the area covered by the raster DTM. As an alternative way of visualising elevation change in the site, cumulative trajectories of elevation change were calculated for a random subset of 10,000 pixels.

To validate the elevation change obtained from LiDAR, we also conducted field measurements of elevation change. We measured elevation change *in situ* at one location within each area (Site A-D and NAT) using 1.5 m metal stakes buried to a depth of 1 m, from which a 50 cm horizontal bar with 10 pins was placed and the distance from the tip of the pin to the bar was measured. Stakes were installed on 14 and 15 December 2014, 14 weeks after the breach, and removed at the end of the study 5-7 March 2017 (Table S1). We found good agreement between LiDAR and ground-based measurements of sedimentation rates (Fig. S2).

### Data analysis: variation in sediment carbon

Variation in sediment carbon composition was assessed as a function of depth using locally weighted polynomial regression (loess function in R), fitted separately for above and below the agricultural soil-new sediment horizon. We assessed whether there was a difference in the percentage carbon in the newly accreted sediment, natural sediment (pooling locations, time points and depths for both) and the pre-restoration soils from the four land uses using Anova with a Tukey HSD post hoc test. Post-restoration samples from at or below the agricultural horizon were not included in this analysis because (1) we were interested in the carbon accumulating after the restoration in the newly accreted sediment and (2) elevated carbon contents were observed due to the burial of remnant agricultural vegetation as opposed to saltmarsh processes. We calculated the mean and standard deviation of TC in newly accreted sediment, and also calculated the mean and standard deviation of the ratio of TOC and TC. We considered it justified to treat the carbon content of new sediment as coming from a single population (i.e. not varying between years) as (1) the TC of new sediment did not vary with depth in cores and (2) there was no difference in the TC of newly accreted sediment (surface sample of sediment in each year) between years (F_1,46_ = 0.369, P = 0.547).

### Data analysis: site-level carbon accumulation

Site-level carbon accumulation was calculated by (1) multiplying mean change in elevation (m, from DTMs) by site area (m^2^) to obtain sediment volume (m^3^), (2) multiplying this by bulk density (t.m^-3^) to obtain sediment mass (t), and (3) multiplying this by sediment carbon content (%/100) to obtain total carbon accumulation (t). This was divided by site area to obtain tC/ha. This calculation was repeated with the additional step of multiplying by the ratio to TOC to TC to estimate site level total organic carbon accumulation.

As each stage in this calculation involves measurements made with error, we used Monte-Carlo resampling to estimate site-level carbon accumulation while propagating errors from each step. If elevation measurement errors were independent for each DTM pixel in each time point then errors largely cancel out. A more conservative approach is to assume that measurement errors apply systematically to a survey. We do the latter, and take mean elevation change between surveys as coming from a normal distribution with a mean equal to the measured change in elevation, and a standard deviation of 0.04 m based on measurements of control points [31]. Bulk density of newly accreted sediment was sampled from a normal distribution with mean 1.11 and SD 0.27 t.m^-3^.Sediment TC was sampled from a normal distribution with mean 4.37% and SD 0.50%, and the ratio of TOC to TC was sampled from a normal distribution with mean 0.53 and SD 0.08. We took 100,000 samples from these distributions to obtain a distribution of carbon accumulation estimates.

### Carbon costs of construction

The carbon cost of constructing the wider 400 ha Steart Marshes complex (comprising the 250 ha managed realignment site and neighbouring areas of freshwater wetland) was estimated using the Environment Agency’s basic carbon calculator (version 3.1.2, dated 2010 (unpublished); since incorporated within the e:Mission Eric carbon planning tool [39]), with the final estimate produced at the end of construction in January 2015. The calculator included estimated greenhouse gas emissions (in carbon dioxide equivalents, CO_2_e) from fuel used for personnel travel, energy use on site within portable accommodation, the emissions embodied in construction materials (considering the weight of material and distance transported to site), and a first order estimate of emissions from machinery fuel usage.

As all the materials for the embankment construction were obtained within the footprint of the site, the principal source of emissions was the fuel consumed by construction machinery moving the material within the site. As such, we have refined the estimate of machinery fuel usage based on the known volume of earthworks undertaken for the whole scheme, where fuel consumption was estimated by considering the work required, fuel burn per hour and productivity per hour. Within the managed realignment site, the amount of material that was excavated and transported was calculated by considering the size of the creek network and the volume of the embankment. The material was excavated using an EC250DL Excavator, transported across the site in a Volvo A25D Articulated Dumper Truck (capacity 10.7 m^3^) and constructed in situ with a D6 Bulldozer and Roller (Table S2). The distance travelled was calculated based on the distance from each section of the embankment to the nearest source of materials, and fuel burn and productivity were obtained from manufacturers and suppliers. Fuel consumption associated with earthworks in other areas of the Steart Marshes complex were based on the volume of earth moved and ground conditions in comparison to the managed realignment site.

## Results

### Sedimentation rates

The change in elevation measured from DTMs was consistent with field measurements of sedimentation (Fig. S2). Comparison of successive DTMs indicated that the net elevation of the site increased over time (Fig. 2), and by September 2018 714,513 m^3^ of sediment had accumulated across the site, with an average depth of 0.292 m (Table 1). There was no clear trend in sedimentation rate with time since breach (regression: slope < 0.001, F_1,8_ = 0.35, P = 0.568), although the most rapid sedimentation was noted immediately following the breach (Table 1). The net elevation of the site increased between most LiDAR surveys. However, in two instances mean elevation decreased between consecutive LiDAR surveys (between June and July 2015, and between March and October 2017), indicating reduction in sediment volume most likely due to dewatering over the summer months.

**Figure 2.**
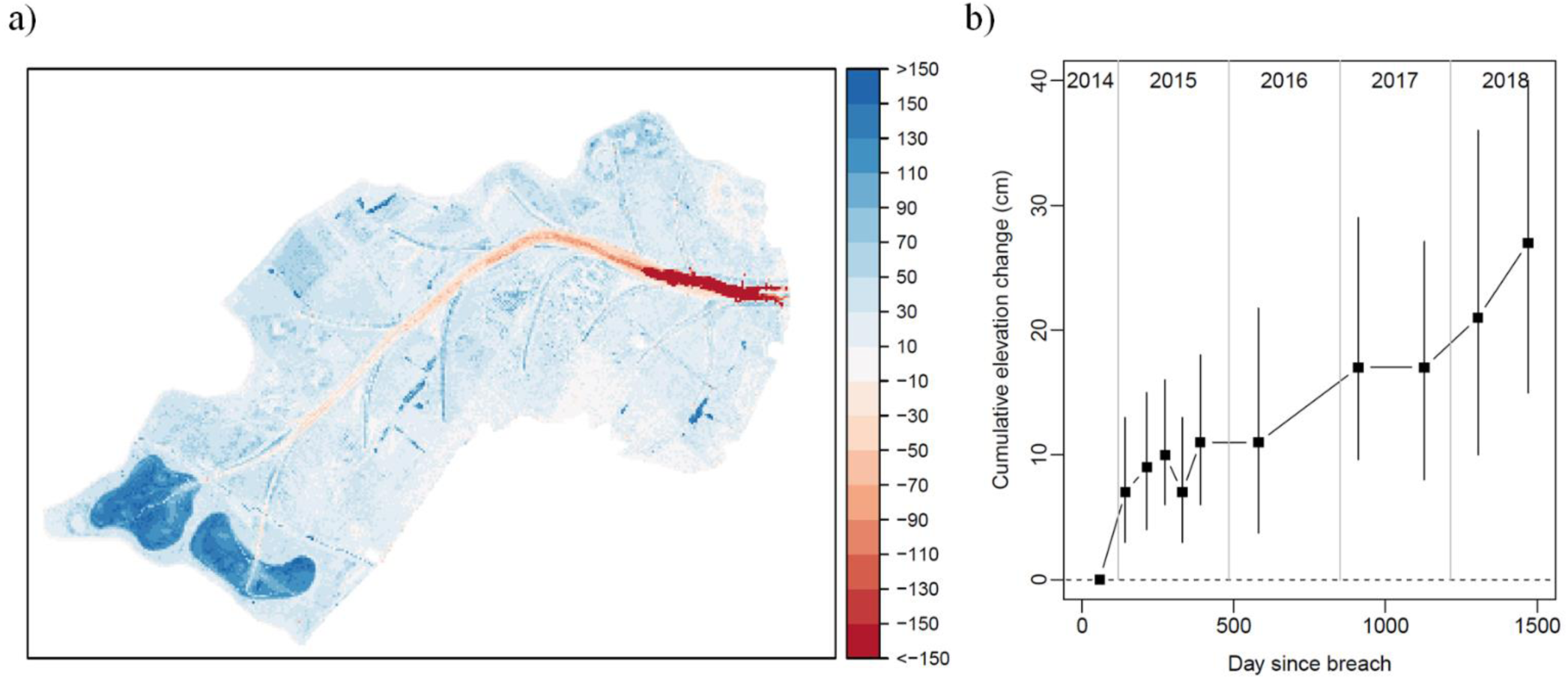
Cumulative sedimentation at Steart Marshes calculated from Lidar DTMs. (a) Change in elevation (cm) between 13/09/2018 (1470 days since breach) and 31/10/2014 (57 days since breach). (b) Cumulative change in elevation over time for individual 50×50 cm pixels. Points show median cumulative change for a random sample of 10,000 pixels. Error bars show the interquartile range for the same sample of pixels.

Within the site, elevation change varied from net accretion of 2.2 m to net erosion of 5.0 m (Fig. 2a), with 92% of DTM pixels experiencing net accretion and 7% experiencing net eroision. Some locations experienced considerable erosion (Fig. S3), expecially in the main creek which deepened progressively in an upstream direction over time (Fig. S4 see [31] for analysis of the main creek profile). Away from the main creek, most locations increased in elevation. This increase was most evident in the excavated pools at the rear of the site, and to a lesser extent in the side creeks (Fig. 2a); 558,648 out of 9,792,179 pixels experienced > 1m of accretion, and 92% of these were located in the two pools.

### Properties of newly accreted sediment

The bulk density of newly accreted sediment ranged from 0.553 to 1.568 t m^-3^ (mean = 1.110 ± 0.267 SD). There were significant differences in TC between pre-restoration soils of different land uses, newly accreted sediments from within the restoration site and sediments from the existing natural saltmarsh (F_5,249_=48.7, p<0.001, Fig. 3). Soils collected prior to restoration from all land uses had significantly lower TC than the newly accreted sediment and the natural saltmarsh sediments, with those from the pre-restoration disturbed (A) and arable (D) areas having the lowest carbon contents (Fig. 3). Sediments from the natural saltmarsh had significantly higher TC (4.72 ± 0.58 %) than the newly accreting sediment on the restoration site (4.37 ± 0.50 %). The ratio of TOC to TC was similar in natural saltmarsh and newly accreting sediment on the restoration site (natural = 0.524, restored = 0.529, Fig. 3), giving a TOC of 2.24 ± 0.33% in newly accreted sediment on the restoration site and 2.44 ± 0.31 % on the natural saltmarsh.

**Figure 3.**
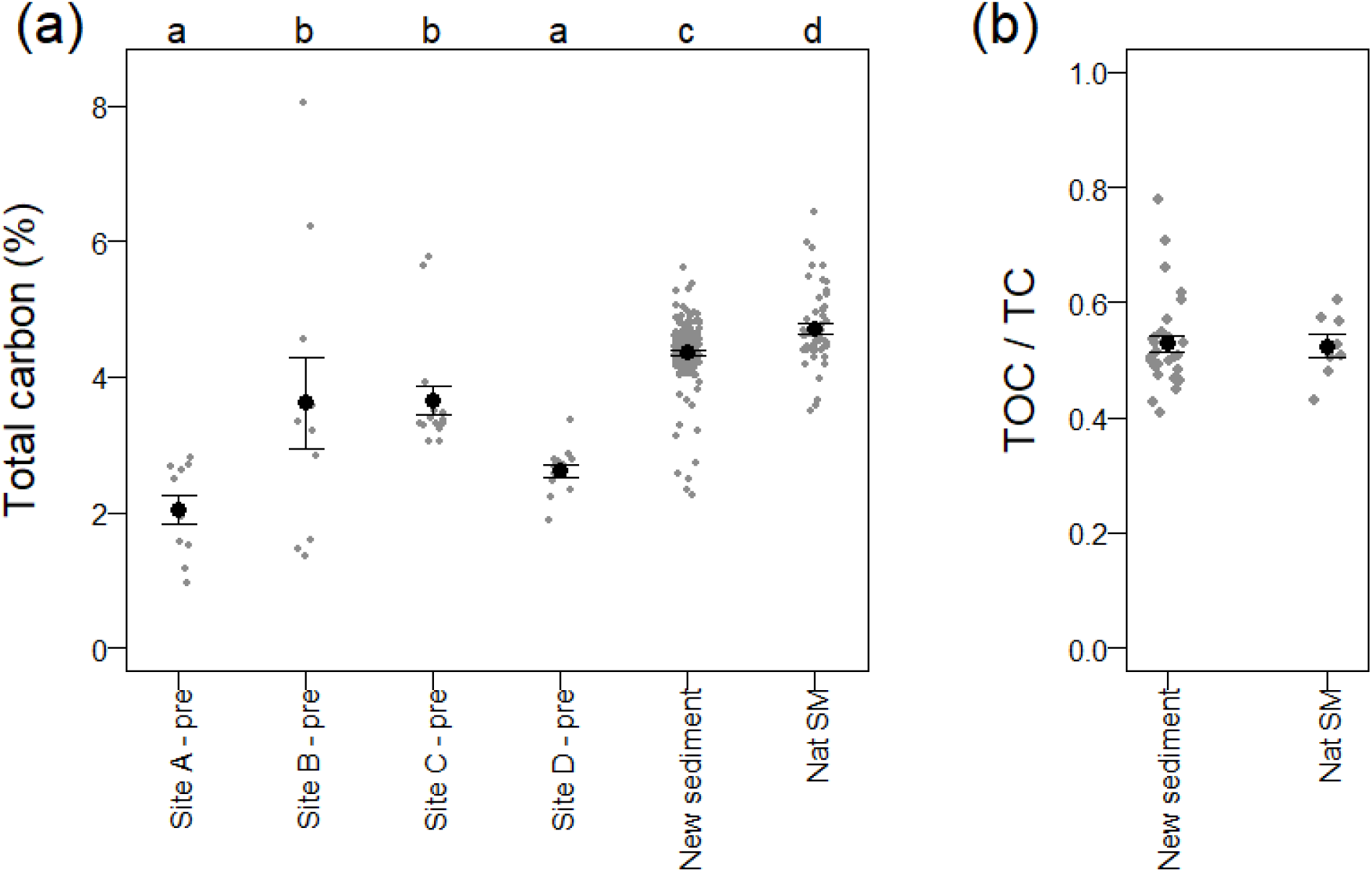
Proportion of total carbon in soil and sediment samples collected from Steart Marshes before and after the restoration of tidal inundation. Soil samples were collected prior to restoration from an area heavily disturbed during construction (site A), an area of pasture (site B), grass ley (site C) and arable (site D). ‘New sediment’ are samples of newly accumulated sediments from the restored site after restoration, with data from all locations and time points pooled. Sediment was also collected from an adjacent natural saltmarsh. Differing letters denote significant differences in the carbon content of sediments between locations (P < 0.05).

There was some spatial variation in the TC of new sediment (significant difference between sampling sites (F_3,44_ = 5.1, P = 0.004)) but no difference between years (F_1,46_ = 0.369, P = 0.547). The TC of newly accreted sediment was consistent with depth (Fig. 4). Some samples taken at the horizon with the underlying agricultural soils had very high carbon content, reflecting the terrestrial vegetation burried by the initial inundations of sediment. Below the horizon, TC was lower than in newly accreted sediment and directly comparable to the pre-breach measurements of the agricultural soils (Fig. 4).

**Figure 4.**
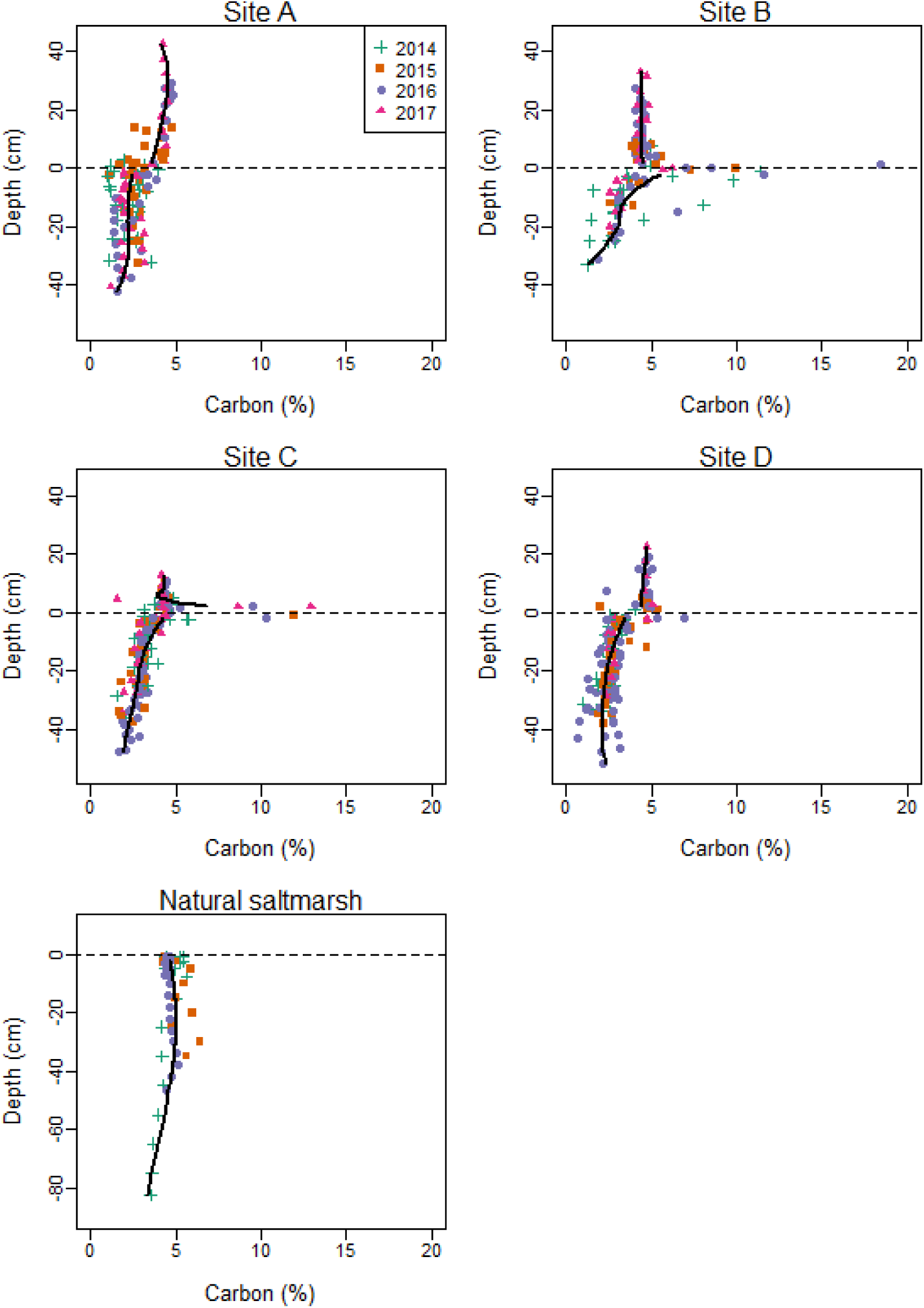
Relationship between soil carbon content and depth. Cores were taken each year at three locations in each starting land-use. Depths are expressed relative to the horizon between agricultural soil and newly deposited sediment, except for natural saltmarsh where depths are from the surface (note difference in y-axis scale for natural saltmarsh). Lines show fits of locally weighted polynomial (loess) models pooling data across locations and years. Loess models have been fit to new sediment (depth > 2 cm) and old sediment (depth < −2 cm) to reduce the effect of vegetation on the horizon.

### Carbon balance

Between 31 October 2014 and 13 September 2018 714,513 m^3^ sediment accumulated on the site. Based on the measured properties of this sediment (mean bulk density of 1.110 ± 0.267 t m^-3^ SD, TC of 4.367 ± 0.499 %) this equates to 34,642 tC (95 % confidence intervals = 16,398 – 57,400) accumulated in sediment, at a rate of 36.6 t C.ha^-1^.yr^-1^ (95% CI = 17.3 – 60.6). Restricting this to TOC (53.0 ± 7.8% of TC) gives 18,343 tC (95% CI = 8090-32402) accumulating at 19.4 tC.ha^-1^.yr^-1^ (95% CI = 8.5-34.2).

We estimated the carbon costs of site construction to compare to the carbon accumulation of the site. In total, 489,422 m^3^ of material were excavated on site, with 411,397 m^3^ used in the construction of the new flood embankments and the remainder used in site landscaping. Moving material across the site resulted in vehicles travelling 69,563 km. Overall, construction of the managed realignment site earthworksrequired 551,012 litres of diesel fuel to be combusted, resulting in 1,477 tCO_2_e (403 tC) being emitted (Table S2). An estimated additional 20% of fuel consumption was assumed for the construction of earthworks in other areas of the Steart Marshes complex, giving total emissions associated with machinery fuel usage of 1,772 tCO_2_e (483 tC).

Combining these figures with the estimated emissions from personnel travel, energy use in portable accomodation, and embodied emissions of construction materials from the Environment Agency carbon calculator, gives estimated total construction emissions of 2,762 tCO_2_e (753 tC). These emissions are equivalent to ∼2% of the of the estimated TC accumulation, or ∼4% of the estimated TOC accumulation in the sediments over the 4 year study period, with a carbon payback period on the order of 1 (TC) to 2 (TOC) months.

## Discussion

We find that Steart Marshes managed realignment has rapidly accumulated carbon since the fronting flood defence embankment was breached, and that this carbon accumulation is two orders of magnitude greater than the carbon costs incurred during site construction. The rate of carbon accumulation at Steart Marshes (TC = 36.6 t C.ha^-1^.yr^-1^, TOC = 19.4 t C.ha^-1^.yr^-1^) is considerably higher than has been found at other sites. In the Bay of Fundy, which like the Severn Estuary is hypertidal, carbon accumulation is lower but within the same order of magnitude at 13.29 t C ha^-1^ yr^-1^ [25], but rates at other sites are an order of magnitude lower than at Steart Marshes. For example, saltmarshes in eastern England were reported to accumulate carbon at a rate of 1.04 t C ha^-1^ yr^-1^ for the first 20 years following creation [24], while a recovering saltmarsh in Australia accumulates at a rate of 0.5 t C ha^-1^ yr^-1^ [40].

The rate of carbon accumulation in a restored saltmarsh is a product of the rate of sediment accumulation and the carbon content of that sediment, and we can look at both these elements to see if Steart Marshes is unusual. Steart Marshes has experienced rapid sediment accumulation since it was breached (mean rate of increase in elevation = 75 mm yr^-1^, Table 1). Similarly high accretion rates have been reported from elsewhere in the Severn Estuary system (short-term accretion rates of 60mm yr^-1^ in young marshes in Bridgwater Bay [41]; around 60mm yr^-1^ in accreting natural marsh in Portishead [42, 43]) and also in the Bay of Fundy (>60 mm yr^-1^ [25]). In comparison, reported sedimentation rates for Tollesbury and Freiston Shore managed realignment sites in eastern England are considerably lower (< 20 mm yr^-1^ [44-46]). A recent meta-analysis has indicated that sediment availability is the dominant control on the vertical accretion of coastal wetland restoration projects, with tidal range, elevation within the tidal frame and sea-level rise explaining a smaller amount of observed variation in the vertical accretion rates of saltmarshes [47]. Hypertidal systems such as the Severn Estuary and the Bay of Fundy are characterised by very high energies and dynamic intertidal sedimentation, where the high suspended sediment load (due to the turbulence created by tidal currents and bores) allows deposition during both flood and ebb tides [48].While the suspended sediment concentrations within the Severn Estuary vary significantly (depending on geographical location, position in the water column, and state of tide), there is a turbidity maxima located in the lower estuary in the vicinity of Bridgewater Bay and the Parrett Estuary (and thus Steart Marshes), with high suspended sediment concentrations typically in the range of 1,000-10,000’s mg/l with values often exceeding 100,000 mg/l [49-52]. Much lower suspended sediment concentrations (∼50-150 mg/l), and thus lower sediment supply, are reported for the Blackwater Estuary (Tollesbury) and The Wash (Freiston Shore) [53, 54].

Sediment carbon content at Steart Marshes is higher than in some managed realignment sites, but is close to the range of values reported elsewhere. Comparison with values from other managed realignments indicates bulk density varies from 0.74 – 1.4 t m^-3^ [24, 55-57] (cf 1.1 in this study) and carbon content varies from 1.8-4.23% (cf TC 4.4% and TOC 2.2% in this study). Combining all combinations of sediment carbon content and accretion rates gives the space of potential carbon accumulation rates in saltmarsh restored by managed realignment (Fig. 5). This indicates that Steart Marshes has high rates of carbon accumulation because it experiences both high rates of accretion and has relatively high sediment carbon content; thus while neither variable is exceptionally high compared with other values reported in the literature, this combination leads to the exceptionally high rates of carbon accumulation. Lower values of either one of these limits carbon accumulation. For example, In natural saltmarshes in China, carbon accumulation rates are low (0.35-3.61 tC ha^-1^ yr^- 1^) despite accretion of 20 mm yr^-1^ because of low sediment carbon densities (< 0.01 g cm^-3^) [58].

**Figure 5.**
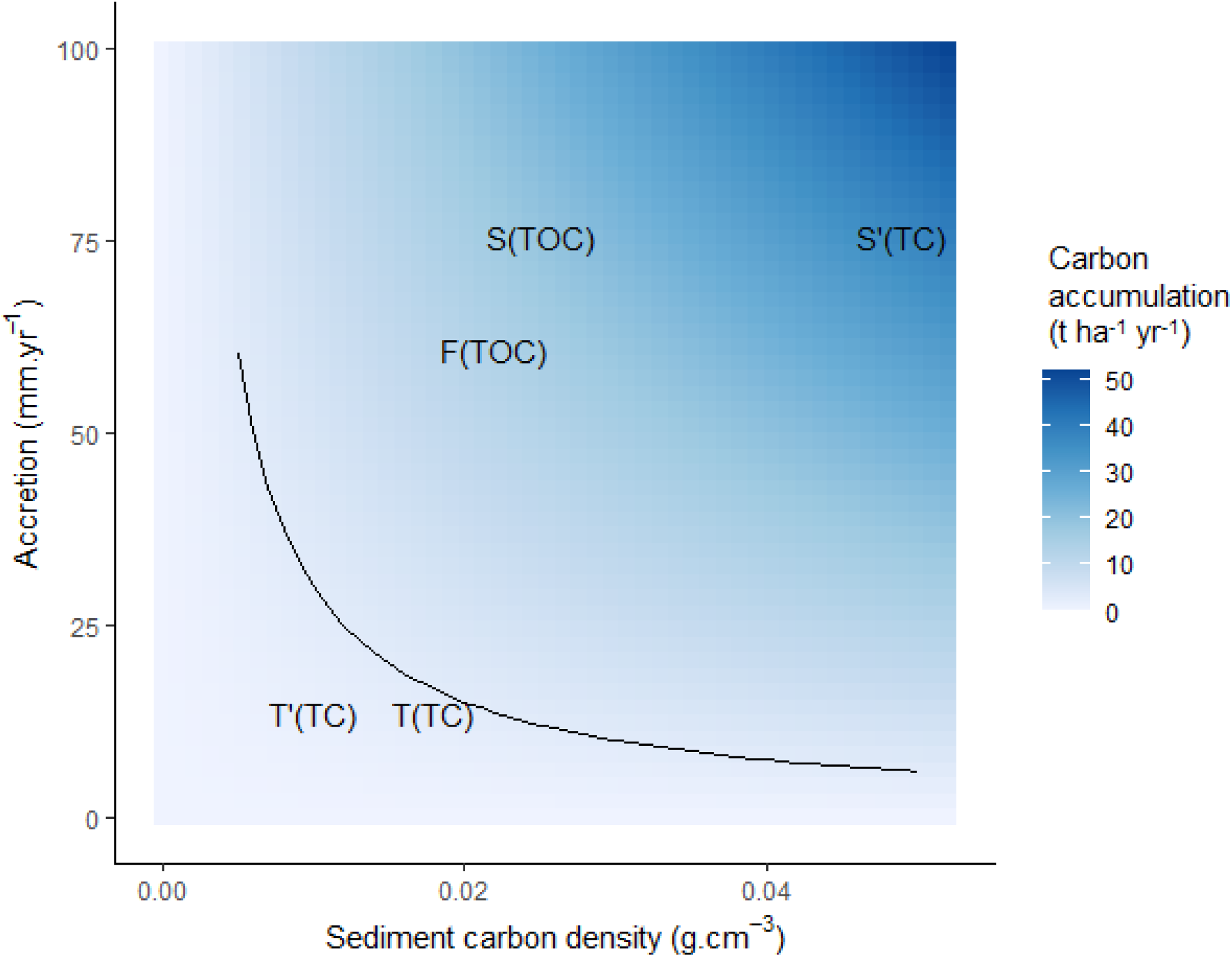
Carbon accumulation potential (tC ha^-1^ yr^-1^) of saltmarsh restored by managed realignment. Values from Steart (S and S’, this study) and published studies at Tollesbury (T [high marsh] and T’ [low marsh] from [45, 69]) and the Bay of Fundy (F, [25]) are shown. TC indicates total carbon sediment carbon density, and TOC indicates total organic carbon sediment carbon density. The solid line deliniates the carbon accumulation rate that would be needed for a site to break even with the per hectare construction carbon costs calculated for Steart in one year. Values to the bottom left of the line fail to break even in a year.

This variability in carbon accumulation rates between sites highlights the need for further work to support large-scale assessments of the carbon sequetration potential of saltmarsh restoration. For example, TOC accumulation rates at Steart Marshes are over 18 times higher (and TC accumulation rates are over 35 times higher) than those used in a recent study to estimate the UK’s carbon sequestration potential [59], while Mossman et al. [28] found ∼13 fold variation in the potential amount of carbon accumulated by restored saltmarshes in the UK based on published estimates of carbon accumulation.

If the high rate of carbon accumulation at Steart Marshes is unusual, is the conclusion that saltmarshes restored by managed realignment rapidly pay off their carbon construction costs applicable to other sites? We can evaluate this by mapping our estimates of construction carbon costs onto the potential carbon accumulation space (Fig. 5). Most combinations of accretion rate and sediment carbon content would pay off construction costs of a 250 ha site within a year, and even a site with low carbon accumulation rates (Tollesbury managed realignment, eastern England) would be close to breaking even with its carbon construction costs over one year.

Our analysis assumes that soil properties (soil carbon, bulk density) come from a single statistical population across the site and over time. However, there were small differences in the carbon content of new sediment across the site. The reasons for this are unclear, but could relate to spatial variation in algal films and vegetation establishment across the site. If the drivers of variation in sediment carbon across the site were known this could be used to scale-up and refine estimates, but this is not currently possible. The bulk density of sediment would be expected to exhibit temporal variation, with lower bulk density (but greater sediment volume) when sediment is waterlogged (e.g. winter, spring), and higher bulk density (but lower sediment volume) when sediment is dry (e.g. summer, early autumn). Our bulk density measurements come from spring and summer, so should capture this temporal variation in bulk density. However, explicitly quantifying temporal variation in bulk density would allow temporal coupling with sediment accumulation data and thus refined quantification of intra-annual variation in carbon accumulation – apparent reductions in carbon stocks over the summer when sediment volume reduced may not occur in reality because of a concurrent increase in sediment bulk density.

### Future changes in carbon accumulation

Although we found the fastest rates of accretion shortly following breaching, we did not find a statistically significant reduction in accretion rates. However, a reduction in accretion rates would be expected as the saltmarsh develops. This is because accretion rates tend to be faster at lower elevations which experience more frequent tidal inundation [60], and as these lower areas increase in elevation they experience fewer inundations, and thus slower accretion. Indeed, space-for-time substitutions indicate that carbon accumulation rates slow over time [24]. It is likely that carbon accumulation rates at Steart Marshes would slow with longer monitoring. Assuming there is sufficient sediment available (as very likely in this case), accretion at managed realignments is expected to occur until the site is a level plane accreting in line with sea-level rise [60]. Natural saltmarsh surrounding the site occurs at elevations of 6.5 m, where if accretion at Steart Marshes stablised at this level this would result in a TC accumulation in excess of 100 ktC (TOC in excess of 50 ktC), of which 33% has currently been accumulated. Even after this point, saltmarshes can continue to accrete with sea-level rise assuming there is sufficent sediment [61], which at 3.7mm yr^-1^ [62] would result in continued TC accumulation of 439 t C yr^-1^ and TOC accumulation of 225 t C yr^-1^ at Steart Marshes assuming no change in sediment carbon content.

### Challenges with determining the carbon budget of a managed realignment

Our results indicate that carbon accumulation at Steart Marshes greatly exceeds construction costs. However, there are a number of uncertanties that, while highly likely not to affect this qualitative conclusion, would need to be considered to refine the quantitive carbon budget (Table 2, [28]).

**Table 2.** Elements that require consideration in the quantification of a full carbon budget of a managed realignment site. The aspects included in this study, the approaches to these that we took, and any implications of these approaches are also given.

| Element | Approach in this study | Rationale of approach and its implications |
| --- | --- | --- |
| Amount of sediment gained and lost within the site | Measured using LiDAR derived DTMs, validated against *in situ* measurements. | Baseline LiDAR is 57 days after breach and ~37 tides covered at least some of the marsh surface between the breach and this LiDAR image, so some post-breach sedimentation will have been missed. This approach has likely underestimated total carbon gained. |
| Changes in intertidal habitat in the wider estuary: the realignment site might cause changes to the sediment dynamics of the wider estuary. Gains and losses of intertidal habitat outside of the site (and the carbon stored in those habitats) that are a direct result of the restoration should be considered | Not considered in this study. | A five year monitoring programme for the scheme found no evidence that the scheme had caused increased erosion in the main estuary channel bed [70]. Near to the scheme, the most significant changes have been associated with the erosion of the exit channel due to the strong flows into and out of the realignment site [31]. Erosion in the exit channel has progressed into the site through the creation of a distinct step, and these changes within the site are captured in our analysis. |
| Carbon in sediment gained | Cores taken at 11 locations in the restored site, average carbon content of new sediment used. | We found total carbon and total organic carbon in new sediment was somewhat lower than in the adjacent natural saltmarsh. The site was in the early stages of restoration (first 4 y). Further development of biotic communities, particularly the vegetation, may increase the carbon content of the sediment at the restored site. Some spatial variation in carbon contents of new sediment around the site. Reasons for this were not clear but understanding this |
|  |  | would allow more spatially refined models of carbon accumulation could be made. |
| Carbon in sediment lost (eroded) | Assumed to be the same as carbon gained. | Carbon content of the soils eroded (e.g. from main creek) is likely to be lower than that in the new sediment because the agricultural soils significantly had lower carbon. However, this could not be quantified because erosion in the main creek was up to 5 m deep. Some of the erosion later in the study would have been of newly accreted sediments (e.g. due to formation of small creeks) and thus of same carbon content as that gained. In total, this approach has likely underestimated total carbon gained. |
| Source of carbon in sediment | Not considered | Burial of *in situ* derived carbon would be a true gain. Carbon from outside the site may have ended up being stored elsewhere in the absence of the site, or may have been oxidised; the extent to which either happens is uncertain. The total carbon accumulation here provides an upper bound for net carbon buried. |
| Plant biomass | Not considered | Belowground biomass contributes to carbon accumulation, and is expected to increase as the site became more vegetated. Similarly, more vegetation creates a source of carbon to be buried. |
| Greenhouse gas fluxes | Not considered | Release of greenhouse gases (e.g. methane) may offset some carbon accumulation. Burden *et al.* [69] found CH<sub>4</sub> and N<sub>2</sub>O fluxes were close to zero on a restored and natural saltmarsh in Essex. However, Adams, Andrew & Jickells [71] suggest gas fluxes |
|  |  | could reduce carbon sequestration on MR sites by 24%. |
| Construction carbon costs | Calculated using estimated fuel use during construction in combination with estimates from the Environment Agency's basic carbon calculator (version 3.1.2) for personnel travel, energy use in portable accommodation, and the embodied emissions associated with construction materials. | Creation of the embankments is very likely to be the greatest construction carbon cost of the managed realignment, and itself is small compared to the carbon gained by the habitat created on the site.<br>All material for the managed realignment part of the site were locally-won material for the embankments from borrow pits on site and was not imported.<br>Since the completion of the Steart project, a more up-to-date Environment Agency carbon estimation tool became available, which considers the carbon of other project stages (operational, decommissioning) to provide the whole-life carbon over a 100-years. The tool currently cannot be adjusted to deal with locally-won embankment material, and was not used here.<br>Some operational carbon cost will occur from the site managers WWT, but has not been included in the calculations. This would include activities associated with site inspections and the maintenance of the embankments (such as grazing by sheep rather than mowing). It is expected to be minimal compared with construction. No decommissioning is anticipated. |
| Prior land use – some changes in land use may result in substantial carbon lost, e.g. loss of trees | Not considered | The site was a mix of arable and pasture prior to restoration and relatively few trees were removed during construction, but this should be considered for future sites. |

Some assumptions, such as assuming the carbon content of sediment lost is the same as sediment gained, are likely to mean our estimate of carbon accumulation is conservative (Table 2). Others, such as not accounting for greenhouse gas emssions following site flooding, will offset some of the carbon accumulation benefit of the site (Table 2).

A particularly challenging element of quantifying the carbon budget of a restored saltmarsh is determining the nature and origin of the carbon accumulated in sediment. This is critical to establish the additionality test needed for carbon codes and offsetting credits (e.g. Verified Carbon Standard Methodology VM0033 [63]), where in simple terms, carbon credits should only be generated through the creation of new net sinks of atmospheric carbon. In the coastal and marine environment, this requires consideration of the relative importance of allocthonous and autochthonous carbon, downstream effects, and both organic carbon and biogenic carbonate inputs. For example, concerns regarding additionality have led to variable treatment of allocthonous carbon, where creating a new apparent store may be depleting supply to an adjacent system (i.e. the carbon would have been stored elsewhere in the absence of the project). Tools such as biomarkers [64] and stable isotopes are being developed to better identify sources of carbon [18], where integrated studies of interconnected blue carbon ecosystems across the land-ocean transect would help address the appropriateness of accounting for allochthonous carbon. While lithogenic carbonates should clearly be excluded from estimates of carbon storage (as they comprise fossil C that has not been in recent contact with the atmosphere), the treatment of biogenic carbonates is more complex. Carbonate production is a source of CO_2_ to the atmosphere, while carbonate dissolution is a sink [65], where the relative balance of these processes and the impact on ecosystem carbon budgets (which also depends on whether the carbonate is imported or produced in-situ, and the role of carbonates in stabilising organic carbon stores) remains a significant uncertainty in blue carbon science [18].

## Conclusions

Our results show that at Steart Marshes fast rates of sediment accumulation and high sediment carbon content combine to result in exceptionally fast carbon accumulation rates. Carbon accumulation at Steart Marshes over the first four years following reinstatement of tidal flow is two orders of magnitude larger than the carbon costs of site construction. Thus qualitatively, it is clear that the creation of the site by managed realignment has delivered benefits for carbon storage and sequestration, and other sites with lower carbon accumulation rates are likely to rapidly pay off their construction carbon debt. However, there are numerous uncertainties that would need to be resolved in order to move to a fully quantitative carbon budget for restored saltmarshes.

## Acknowledgements

We thank the Wildfowl and Wetlands Trust, particularly Alys Laver and Tim McGrath, for access to the site and their ongoing enthusiasm and support. We thank Grace Biddle, Colin Hill and David McKendry for their work in the laboratory. This study uses data from UK National Tide Gauge Network, owned and operated by the Environment Agency, and provided by the British Oceanographic Data Centre.

## SUPPLEMENTARY INFORMATION

**Table S1.** Field sampling dates and information. Samples highlighted in bold are those selected for the quantification of total organic carbon. Access issues prevented sampling at some locations in March 2015 and September 2016. We assessed the consequence of the additional uneven sampling by removing samples from these two time periods and recalculating the mean carbon content in newly accreted sediment. The value differed by less than 1% of the original value (i.e. 4.367% vs 4.372%), so we retain all samples in the data presented in the manuscript.

| Date of sampling | Days after restoration | Cores sampled | Notes |
| --- | --- | --- | --- |
| 28 August 2014 | -7 | Site A1, A2<br>Site B1, B2<br>Site C1, C2, C3<br>Site D1, D2, D3 | Pre-restoration sampling. Natural marsh not sampled.<br>A3 not sampled as it was undergoing earthworks. |
| 14-15 December 2014 | +101 | Site A1, A2, A3<br>Site B1, B2<br>Site C1, C2, C3<br>Site D1, D2, D3<br>NAT | Installation of sedimentation pins |
| 16-17 March 2015 | +193 | Site A2<br>Site C1, C2, C3<br>Site D1, D2, D3<br>NAT | Access issues prevented sampling at Site B, A1 and A3. |
| 25-28 June 2015 | +294 | Site A1, A2, A3<br>Site B1, B2<br>Site C1, C2, C3<br>Site D1, D2, D3<br>NAT |  |
| 14-15 April 2016 | +588 | Site A1, A2, A3<br>Site B1, B2<br>Site C1, C2, C3<br>Site D1, D2, D3<br>**NAT** |  |
| 27-28 September 2016 | +754 | Site B1, B2<br>Site C1, C2<br>Site D1, D2, D3 | Access issues prevented sampling at Site A and C3. |
| 5-7 March 2017 | +913 | Site A1, A2, **A3**<br>Site B1, **B2**<br>Site C1, C2, **C3**<br>Site D1, **D2**, D3 | Sedimentation pin data collected |

**Table S2.** Summary of the fuel (diesel) consumption and tCO_2_e emitted by machinery in the construction of Steart Marshes earthworks.

| Equipment/Component | Average fuel burn (l.hr<sup>-1</sup>) | Productivity | Fuel use (l) | t CO<sub>2</sub> | tC |
| --- | --- | --- | --- | --- | --- |
| Excavator (EC250DL) | 20.8<sup>2</sup> | 80m<sup>3</sup>/hr<sup>6</sup> | 127,255 | 341.0 | 93.0 |
| A25D ADT | 30<sup>1</sup> | 10 km/hr<sup>5</sup> | 208,689 | 559.3 | 152.5 |
| D6 Bulldozer | 19<sup>3</sup> | 45.7m<sup>3</sup>/hr<sup>7</sup> | 203,488 | 545.3 | 148.7 |
| Roller | 1.9<sup>4</sup> |  | 11,580 | 31.0 | 8.5 |
| Managed Realignment Earthworks |  |  | 551,012 | 1,476.7 | 402.7 |
| Other Earthworks |  |  | 110,202 | 295.3 | 80.5 |
| **Total Earthworks** |  |  | **661,214** | **1,772** | **483** |

<sup>1</sup>https://www.equipmentworld.com/owning-and-operating-costs-5; <sup>2</sup>https://www.volvoc.com/en-us/government/services/promotions/fuel-efficiency-guarantee; <sup>3</sup>https://www.constructionequipmentguide.com/industry-begins-grappling-with-rising-fuel-costs/7008; <sup>4</sup> https://www.equipmentworld.com/owning-and-operating-costs-6/; <sup>5</sup> Estimated from average speed on site (obtained from the contractor); <sup>6</sup> Hydraulic Excavator with a 0.5 m<sup>3</sup> bucket excavating clay/chalk, https://www.methvin.org/construction-production-rates/excavation/bulk-excavation; <sup>7</sup> 1.2 m<sup>3</sup> bucket, reach of 50-200 m<sup>2</sup>, Boulder Clay https://www.methvin.org/construction-production-rates/excavation/bulk-excavation

**Figure S1.**
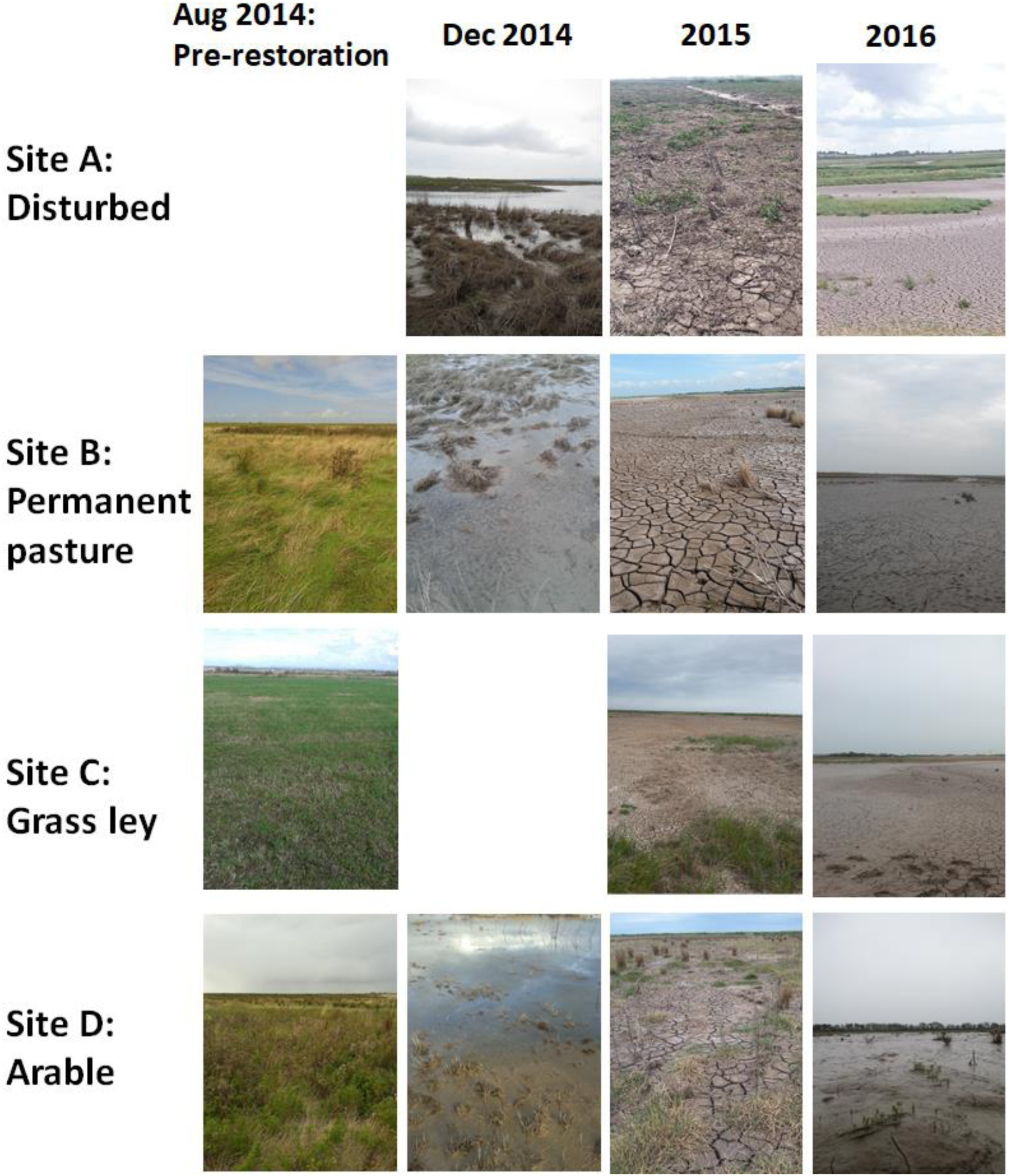
Photographs of sampling areas (Sites A-D).

**Figure S2.**
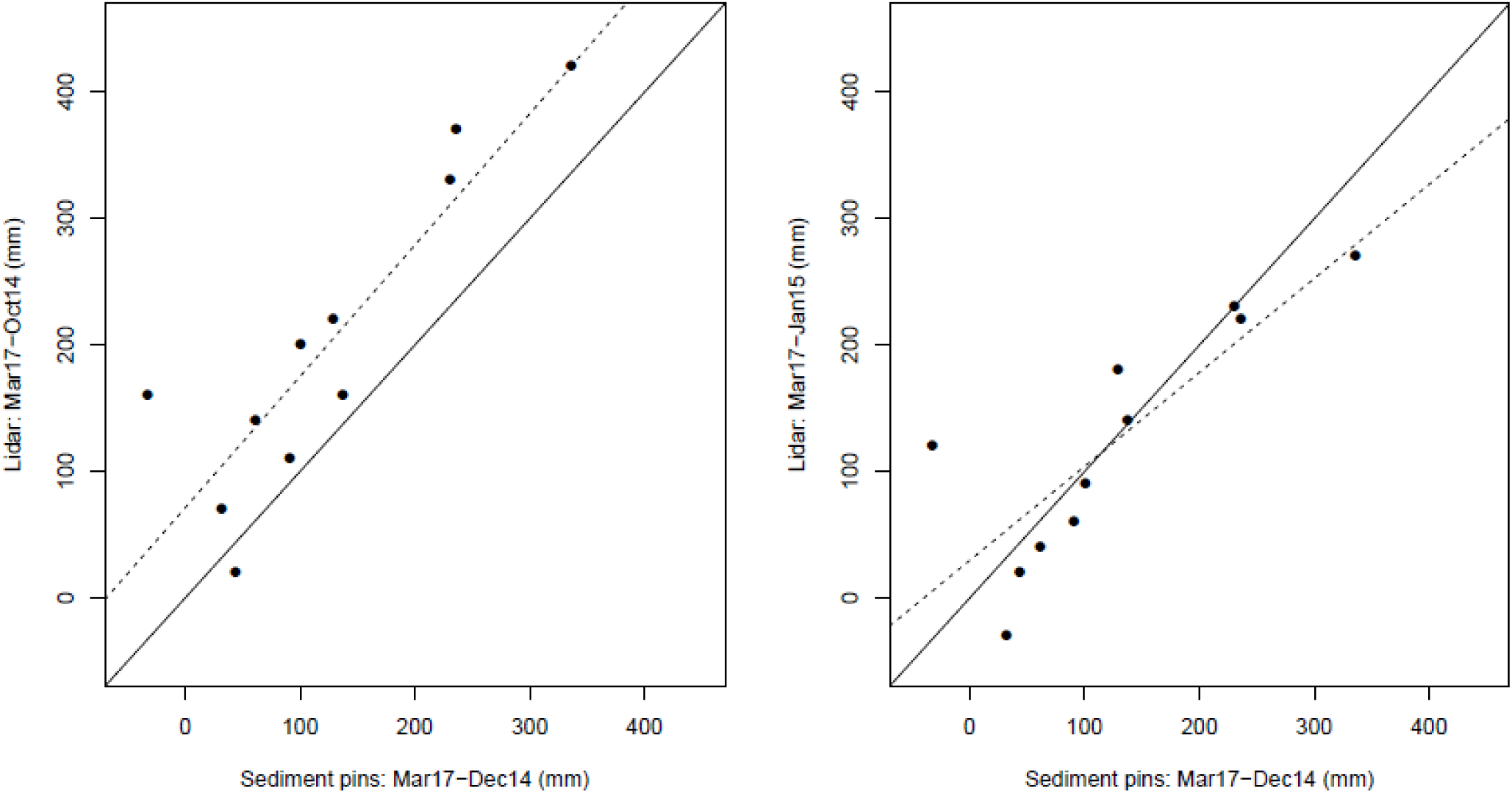
Relationship between elevation change measured with LiDAR derived-DTMs and in situ measurements with pins. In situ measured data (x axis) show difference in elevation between December 2014 (3 months after restoration) and March 2017. Left: Compares in situ data to elevation changes derived from LiDAR data taken in October 2014 and March 2017, and Right compares elevation changes between January 2015 and March 2017. No LiDAR images are available for December 2014. Solid lines show a 1:1 relationship and the dashed lines show the actual relationship (linear regression) between DTM-derived and in situ measurements (dash lines Left: R^2^ = 0.775, P <0.001; Right R^2^= 0.686, P = 0.002). LiDAR measurements are strongly related to in situ measurements and are not systematically biased when sampling periods are more closely matched (i.e. Right).

**Figure S3.**
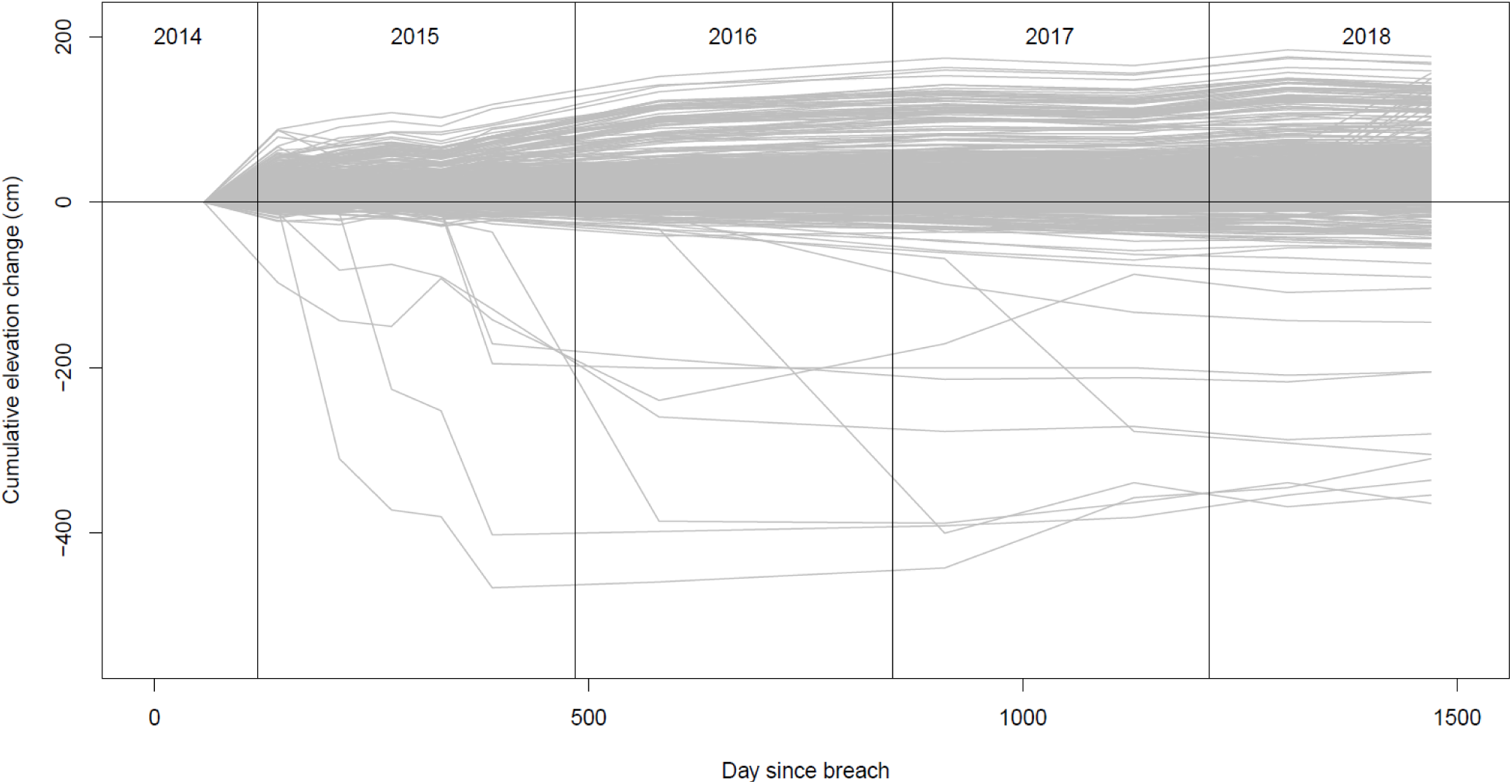
Cumulative elevation change trajectories of a sample of 1000 DTM pixels.

**Figure S4.**
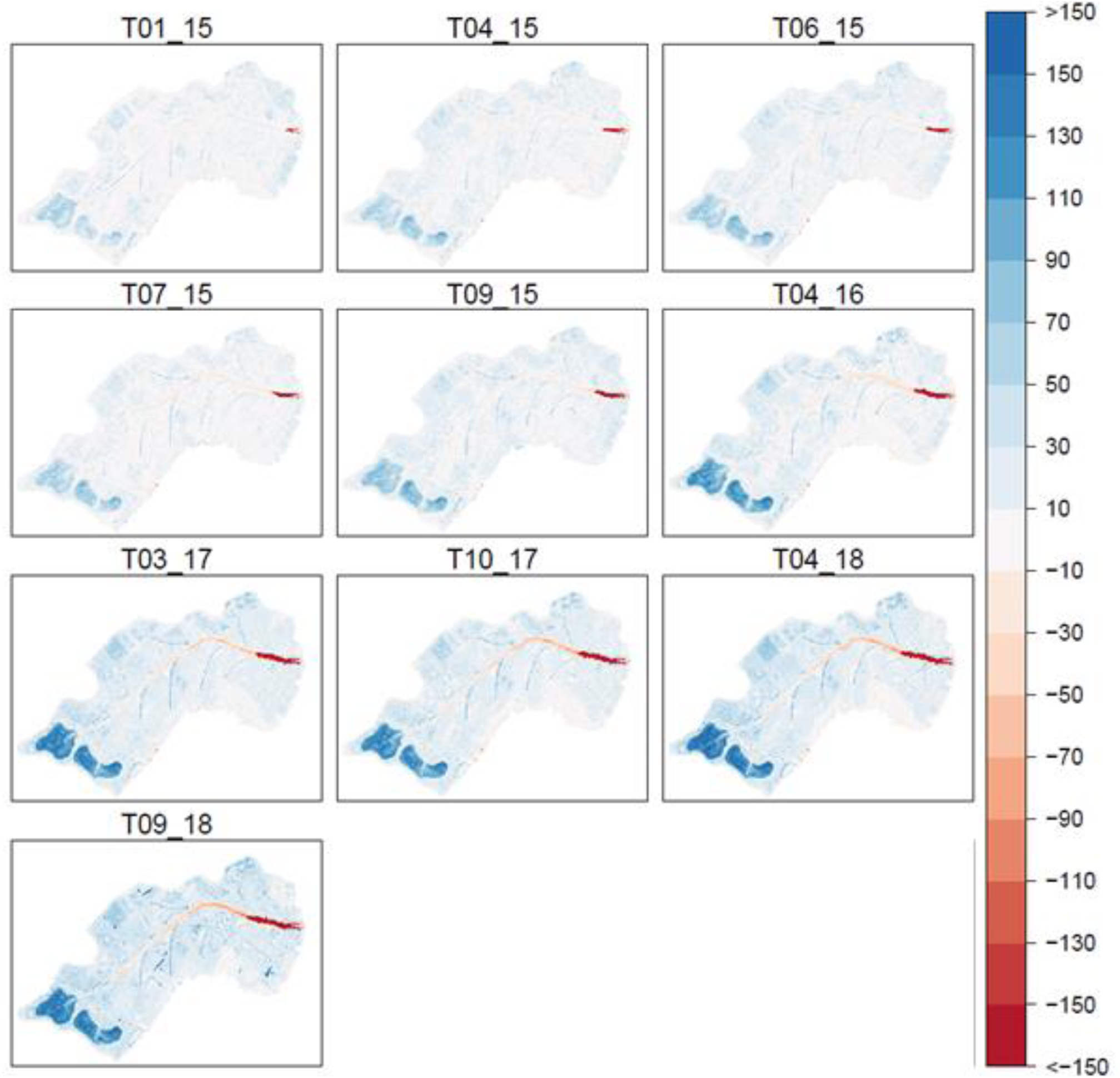
Cumulative change in elevation for each LiDAR survey.

## References

1. Friedlingstein P, O’Sullivan M, Jones MW, Andrew RM, Hauck J, Olsen A, et al. Global Carbon Budget 2020. Earth Syst Sci Data. 2020;12(4):3269–340. doi: 10.5194/essd-12-3269-2020.

2. Guo LB, Gifford RM. Soil carbon stocks and land use change: a meta analysis. Global Change Biology. 2002;8(4):345–60. doi: https://doi.org/10.1046/j.1354-1013.2002.00486.x.

3. Sullivan MJP, Lewis SL, Affum-Baffoe K, Castilho C, Costa F, Sanchez AC, et al. Long-term thermal sensitivity of Earth’s tropical forests. Science. 2020;368(6493):869–74.

4. McLeod E, Chmura GL, Bouillon S, Salm R, Björk M, Duarte CM, et al. A blueprint for blue carbon: toward an improved understanding of the role of vegetated coastal habitats in sequestering CO2. Frontiers in Ecology and the Environment. 2011;9(10):552–60. doi: https://doi.org/10.1890/110004.

5. McOwen CJ, Weatherdon LV, Van Bochove J-W, Sullivan E, Blyth S, Zockler C, et al. A global map of saltmarshes. Biodiversity data journal. 2017;(5).

6. Ouyang X, Lee SY. Updated estimates of carbon accumulation rates in coastal marsh sediments. Biogeosciences. 2014;11(18):5057–71.

7. Barbier EB, Hacker SD, Kennedy C, Koch EW, Stier AC, Silliman BR. The value of estuarine and coastal ecosystem services. Ecological Monographs. 2011;81(2):169–93. doi: 10.1890/10-1510.1. PubMed PMID: WOS:000290707600001.

8. Murray NJ, Clemens RS, Phinn SR, Possingham HP, Fuller RA. Tracking the rapid loss of tidal wetlands in the Yellow Sea. Frontiers in Ecology and the Environment. 2014;12(5):267–72. doi: 10.1890/130260.

9. Duarte CM, Dennison WC, Orth RJW, Carruthers TJB. The charisma of coastal ecosystems: addressing the imbalance. Estuaries and coasts. 2008;31(2):233–8.

10. Bull JW, Milner-Gulland EJ. Choosing prevention or cure when mitigating biodiversity loss: Trade-offs under ‘no net loss’ policies. Journal of Applied Ecology. 2020;57(2):354–66.

11. Li S, Xie T, Pennings SC, Wang Y, Craft C, Hu M. A comparison of coastal habitat restoration projects in China and the United States. Scientific Reports. 2019;9(1):14388. doi: 10.1038/s41598-019-50930-6.

12. Stewart-Sinclair PJ, Purandare J, Bayraktarov E, Waltham N, Reeves S, Statton J, et al. Blue Restoration – Building Confidence and Overcoming Barriers. Frontiers in Marine Science. 2020;7:748.

13. Salzman J, Bennett G, Carroll N, Goldstein A, Jenkins M. The global status and trends of Payments for Ecosystem Services. Nature Sustainability. 2018;1(3):136–44.

14. Vieira da Silva L, Everard M, Shore RG. Ecosystem services assessment at Steart Peninsula, Somerset, UK. Ecosystem Services. 2014;10:19–34. doi: https://doi.org/10.1016/j.ecoser.2014.07.008.

15. Serrano O, Lovelock CE, B. Atwood T, Macreadie PI, Canto R, Phinn S, et al. Australian vegetated coastal ecosystems as global hotspots for climate change mitigation. Nature Communications. 2019;10(1):4313. doi: 10.1038/s41467-019-12176-8.

16. Needelman BA, Emmer IM, Emmett-Mattox S, Crooks S, Megonigal JP, Myers D, et al. The Science and Policy of the Verified Carbon Standard Methodology for Tidal Wetland and Seagrass Restoration. Estuaries and Coasts. 2018;41(8):2159–71. doi: 10.1007/s12237-018-0429-0.

17. Wedding LM, Moritsch M, Verutes G, Arkema K, Hartge E, Reiblich J, et al. Incorporating blue carbon sequestration benefits into sub-national climate policies. Global Environmental Change. 2021:102206. doi: https://doi.org/10.1016/j.gloenvcha.2020.102206.

18. Macreadie PI, Anton A, Raven JA, Beaumont N, Connolly RM, Friess DA, et al. The future of Blue Carbon science. Nature Communications. 2019;10(1):3998. doi: 10.1038/s41467-019-11693-w.

19. Lawrence PJ, Smith GR, Sullivan MJ, Mossman HL. Restored saltmarshes lack the topographic diversity found in natural habitat. Ecological engineering. 2018;115:58–66.

20. Mossman HL, Davy AJ, Grant A. Does managed coastal realignment create saltmarshes with ‘equivalent biological characteristics’ to natural reference sites? Journal of Applied Ecology. 2012;49(6):1446–56. doi: 10.1111/j.1365-2664.2012.02198.x.

21. Moreno-Mateos D, Power ME, Comín FA, Yockteng R. Structural and Functional Loss in Restored Wetland Ecosystems. PLOS Biology. 2012;10(1):e1001247. doi: 10.1371/journal.pbio.1001247.

22. Moritsch MM, Young M, Carnell P, Macreadie PI, Lovelock C, Nicholson E, et al. Estimating blue carbon sequestration under coastal management scenarios. Science of The Total Environment. 2021;777:145962. doi: https://doi.org/10.1016/j.scitotenv.2021.145962.

23. MacDonald MA, de Ruyck C, Field RH, Bedford A, Bradbury RB. Benefits of coastal managed realignment for society: Evidence from ecosystem service assessments in two UK regions. Estuarine, Coastal and Shelf Science. 2020;244:105609. doi: https://doi.org/10.1016/j.ecss.2017.09.007.

24. Burden A, Garbutt A, Evans CD. Effect of restoration on saltmarsh carbon accumulation in Eastern England. Biology Letters. 2019;15(1):20180773.

25. Wollenberg JT, Ollerhead J, Chmura GL. Rapid carbon accumulation following managed realignment on the Bay of Fundy. PLOS ONE. 2018;13(3):e0193930. doi: 10.1371/journal.pone.0193930.

26. Hoogsteen MJJ, Lantinga EA, Bakker EJ, Groot JCJ, Tittonell PA. Estimating soil organic carbon through loss on ignition: effects of ignition conditions and structural water loss. European Journal of Soil Science. 2015;66(2):320–8. doi: https://doi.org/10.1111/ejss.12224.

27. The Greenhouse Gas Protocol. The GHG Protocol for Project Accounting. World Business Council for Sustainable Development and World Resources Institute; 2005.

28. Mossman HL, Sullivan MJP, Dunk RM, Rae S, Sparkes RT, Pontee, N. Created coastal wetlands as carbon stores: potential challenges and opportunities. In: Humphreys J, Little S, editors. Challenges in Estuarine and Coastal Science: Estuarine and Coastal Sciences Association 50th Anniversary Volume. UK: Pelagic Publishing; 2021.

29. Scott J, Pontee N, McGrath T, Cox R, Philips M. Delivering Large Habitat Restoration Schemes: Lessons from the Steart Coastal Management Project. Coastal Management: Changing coast, changing climate, changing minds: ICE Publishing; 2016. p. 663–74.

30. British Geological Society. Geology of Britain [cited 2021 21 April]. Available from: https://mapapps.bgs.ac.uk/geologyofbritain/home.html.

31. Pontee N, Serato B. Nearfield erosion at the steart marshes (UK) managed realignment scheme following opening. Ocean & Coastal Management. 2019;172:64–81.

32. UK Hydrographic Office. Admiralty Tide Tables Volume 1: United Kingdom and Ireland (Including European Channel Ports). Taunton, UK: The United Kingdom Hydrographic Office; 2010.

33. Pontee NI. Impact of managed realignment design on estuarine water levels. Proceedings of the Institution of Civil Engineers - Maritime Engineering. 2015;168(2):48–61.

34. Rowell DL. Soil science: Methods & applications: Routledge; 2014.

35. Sparkes RB, Lin I-T, Hovius N, Galy A, Liu JT, Xu X, et al. Redistribution of multi-phase particulate organic carbon in a marine shelf and canyon system during an exceptional river flood: Effects of Typhoon Morakot on the Gaoping River–Canyon system. Marine Geology. 2015;363:191–201. doi: https://doi.org/10.1016/j.margeo.2015.02.013.

36. Defra. LiDAR Composite DTM - 0.5 m. Open Government Licence v3.0 https://environment.data.gov.uk/DefraDataDownload/?Mode=survey2020 [25 January 2021].

37. R Development Core Team. R: A language and environment for statistical computing. 3.5.0 ed. Vienna: R Foundation for Statistical Computing; 2018.

38. Hijmans RJ. raster: Geographic Data Analysis and Modeling. R package version 3.3-6. 2020.

39. Environment Agency. Eric carbon planning tool training package: https://www.ericenvironmentagency.co.uk/story_html5.html?lms=1 [cited 2021 08/10/2021].

40. Gulliver A, Carnell PE, Trevathan-Tackett SM, Duarte de Paula Costa M, Masqué P, Macreadie PI. Estimating the Potential Blue Carbon Gains From Tidal Marsh Rehabilitation: A Case Study From South Eastern Australia. Frontiers in Marine Science. 2020;7:403.

41. Ranwell DS. Spartina salt marshes in southern England: II. Rate and seasonal pattern of sediment accretion. The Journal of Ecology. 1964:79–94.

42. Allen JRL, Duffy MJ. Medium-term sedimentation on high intertidal mudflats and salt marshes in the Severn Estuary, SW Britain: the role of wind and tide. Marine Geology. 1998;150(1-4):1–27.

43. Allen JRL, Duffy MJ. Temporal and spatial depositional patterns in the Severn Estuary, southwestern Britain: intertidal studies at spring–neap and seasonal scales, 1991–1993. Marine Geology. 1998;146(1-4):147–71.

44. Brown SL, Pinder A, Scott L, Bass J, Rispin E, Brown S, et al. Wash Banks Flood Defence Scheme Freiston Environmental Monitoring 2002-2006. Report to Environment Agency, Peterborough. Centre for Ecology and Hydrology, Dorset, UK: 2007.

45. Garbutt A. Bed level change within the Tollesbury managed realignment site, Blackwater estuary, Essex, UK between 1995 and 2007. NERC Environmental Information Data Centre; 2018.

46. Spencer T, Friess DA, Möller I, Brown SL, Garbutt RA, French JR. Surface elevation change in natural and re-created intertidal habitats, eastern England, UK, with particular reference to Freiston Shore. Wetlands Ecology and Management. 2012;20(1):9–33. doi: 10.1007/s11273-011-9238-y.

47. Liu Z, Fagherazzi S, Cui B. Success of coastal wetlands restoration is driven by sediment availability. Communications Earth & Environment. 2021;2(1):1–9.

48. Archer AW. World’s highest tides: Hypertidal coastal systems in North America, South America and Europe. Sedimentary Geology. 2013;284-285:1–25. doi: https://doi.org/10.1016/j.sedgeo.2012.12.007.

49. Thorn MFC, Burt TN. Sediments and metal pollutants in a turbid tidal estuary. Canadian Journal of Fisheries and Aquatic Sciences. 1983;40(S1):s207–s15.

50. Mantz PA, Wakeling HL. Aspects of sediment movement near to Bridgwater Bay bar, Bristol Channel. Proceedings of the Institution of Civil Engineers. 1982;73(1):1–23.

51. Darbyshire EJ, West JR. Turbulence and cohesive sediment transport in the Parrett estuary. Turbulence: Perspectives on Flow and Sediment Transport Wiley, Chichester. 1993:215–47.

52. Manning AJ, Langston WJ, Jonas PJC. A review of sediment dynamics in the Severn Estuary: Influence of flocculation. Marine Pollution Bulletin. 2010;61(1):37–51. doi: https://doi.org/10.1016/j.marpolbul.2009.12.012.

53. French JR. Numerical simulation of vertical marsh growth and adjustment to accelerated sea-level rise, North Norfolk, U.K. Earth Surface Processes and Landforms. 1993;18(1):63–81. doi: https://doi.org/10.1002/esp.3290180105.

54. Spearman J. The development of a tool for examining the morphological evolution of managed realignment sites. Continental Shelf Research. 2011;31(10):S199–S210.

55. Clapp J. Managed realignment in the Humber estuary: factors influencing sedimentation. 2009. Unpublished PhD thesis, University of Hull.

56. Spencer KL, Carr SJ, Diggens LM, Tempest JA, Morris MA, Harvey GL. The impact of pre-restoration land-use and disturbance on sediment structure, hydrology and the sediment geochemical environment in restored saltmarshes.Science of the Total Environment.2017;587:47–58.

57. Blackwell MSA, Yamulki S, Bol R. Nitrous oxide production and denitrification rates in estuarine intertidal saltmarsh and managed realignment zones. Estuarine, Coastal and Shelf Science. 2010;87(4):591–600.

58. Chen J, Wang D, Li Y, Yu Z, Chen S, Hou X, et al. The carbon stock and sequestration rate in tidal flats from coastal China. Global Biogeochemical Cycles. 2020;34(11):e2020GB006772.

59. Bradfer-Lawrence T, Finch T, Bradbury RB, Buchanan GM, Midgley A, Field RH. The potential contribution of terrestrial nature-based solutions to a national ‘net zero’ climate target. Journal of Applied Ecology. 2021;n/a(n/a). doi: https://doi.org/10.1111/1365-2664.14003.

60. Pontee N. Accounting for siltation in the design of intertidal creation schemes. Ocean & coastal management. 2014;88:8–12.

61. Schuerch M, Spencer T, Temmerman S, Kirwan ML, Wolff C, Lincke D, et al. Future response of global coastal wetlands to sea-level rise. Nature. 2018;561(7722):231–4. doi: 10.1038/s41586-018-0476-5.

62. Met Office. UKCP09: Gridded observation data sets. 2009.

63. Emmer I, Needelman B, Emmett-Mattox S, Crooks S, Megonigal P, Myers D, et al. VM0033 Methodology for tidal wetland and seagrass restoration. Version 1.0. Verra. Verified Carbon Standard, 2015.

64. Bischoff J, Sparkes RB, Doğrul Selver A, Spencer RGM, Gustafsson Ö, Semiletov IP, et al. Source, transport and fate of soil organic matter inferred from microbial biomarker lipids on the East Siberian Arctic Shelf. Biogeosciences. 2016;13(17):4899–914.

65. Saderne V, Geraldi NR, Macreadie PI, Maher DT, Middelburg JJ, Serrano O, et al. Role of carbonate burial in Blue Carbon budgets. Nature communications. 2019;10(1):1–9.

66. Centre for Ecology and Hydrology. Land Cover Map 2007 [SHAPE geospatial data], Scale 1:250000. Updated: 18 July 2008.: EDINA Environment Digimap Service, <https://digimap.edina.ac.uk> 2007.

67. Environment Agency. Steart Coastal Management Project Environmental Statement: Report produced by Halcrow for the Environment Agency. Bristol, UK: Environment Agency; 2011. p. 178pp

68. Getmapping. High Resolution (25cm) Vertical Aerial Imagery [JPG geospatial data], Scale 1:500, Updated: 25 October 2014. EDINA Aerial Digimap Service, https://digimap.edina.ac.uk; 2014.

69. Burden A, Garbutt RA, Evans CD, Jones DL, Cooper DM. Carbon sequestration and biogeochemical cycling in a saltmarsh subject to coastal managed realignment. Estuarine, Coastal and Shelf Science. 2013;120:12–20.

70. Jacobs. Final Far Field Effect & Channel Exit: Review and summary – Note 6. Report prepared for the Environment Agency by Jacobs. 2019. p. 39pp.

71. Adams CA, Andrews JE, Jickells T. Nitrous oxide and methane fluxes vs. carbon, nitrogen and phosphorous burial in new intertidal and saltmarsh sediments. Science of the Total Environment. 2012;434:240–51.

